# Generative latent diffusion language modeling yields anti-infective synthetic peptides

**DOI:** 10.1101/2025.01.31.636003

**Authors:** Marcelo D. T. Torres, Tianlai Chen, Fangping Wan, Pranam Chatterjee, Cesar de la Fuente-Nunez

**Author notes:** These authors contributed equally to this work (Marcelo D. T. Torres, Tianlai Chen).

## Abstract

Generative artificial intelligence (AI) offers a powerful avenue for peptide design, yet this process remains challenging due to the vast sequence space, complex structure–activity relationships, and the need to balance antimicrobial potency with low toxicity. Traditional approaches often rely on trial-and-error screening and fail to efficiently navigate the immense diversity of potential sequences. Here, we introduce AMP-Diffusion, a novel latent diffusion model fine-tuned on antimicrobial peptide (AMP) sequences using embeddings from protein language models. By systematically exploring sequence space, AMP-Diffusion enables the rapid discovery of promising antibiotic candidates. We generated 50,000 candidate sequences, which were subsequently filtered and ranked using our APEX predictor model. From these, 46 top candidates were synthesized and experimentally validated. The resulting AMP-Diffusion peptides demonstrated broad-spectrum antibacterial activity, targeting clinically relevant pathogens—including multidrug-resistant strains—while exhibiting low cytotoxicity in human cell assays. Mechanistic studies revealed bacterial killing via membrane permeabilization and depolarization, and the peptides showed favorable physicochemical profiles. In preclinical mouse models of infection, lead peptides effectively reduced bacterial burdens, displaying efficacy comparable to polymyxin B and levofloxacin, with no detectable adverse effects. This study highlights the potential of AMP-Diffusion as a robust generative platform for designing novel antibiotics and bioactive peptides, offering a promising strategy to address the escalating challenge of antimicrobial resistance.

## Introduction

The field of generative artificial intelligence (AI) has experienced significant advancements, with denoising diffusion models emerging as effective tools for diverse tasks in computer vision, natural language processing, and more recently, protein design^1^. By progressively denoising data from a Gaussian noise distribution back to the original signal, diffusion models enable the exploration of innovative strategies for designing proteins with specific structural and functional characteristics.

In protein design, diffusion-based approaches such as RFDiffusion have successfully engineered protein monomers, binders, and symmetric oligomers, leveraging their ability to explore vast conformational spaces^2^. Similarly, graph-based diffusion models have enabled progress in specialized areas such as antibody design and protein-ligand docking^3,4^. By leveraging graph representations of protein structures and interactions, these models refine and optimize molecular configurations, leading to antibodies with improved binding affinities or ligands displaying enhanced docking properties.

Alongside diffusion-based methods, protein language models (pLMs) have emerged as powerful tools for protein sequence analysis and design. Models such as ESM-2^5^, ProtT5^6^, and others are pre-trained on large-scale datasets comprising millions of natural protein sequences. Through transformer architectures, they capture fundamental physicochemical and functional properties directly from the sequences. This capability extends to *de novo* protein design, where pLMs can generate entirely new sequences that retain functionality and bioactivity. For example, ProtGPT2^7^ and ProGen^8^ are pre-trained in an autoregressive manner, while EvoDiff employs diffusion to support both *de novo* sequence generation and scaffolding^9^.

Antimicrobial peptides (AMPs) offer a promising alternative to conventional antibiotics due to their multifaceted mechanisms of action, including membrane disruption, metabolic inhibition, and immune modulation^10–12^. This versatility makes them potent candidates for countering antimicrobial resistance (AMR)^13–15^. Despite their potential, AMP discovery remains challenging. Machine learning (ML) and deep learning (DL) have become indispensable tools in AMP discovery and generation, accelerating research by identifying functional patterns in large datasets^16–26^.

Given the rising threat of AMR, we hypothesized that integrating representation and generative language modeling would facilitate robust AMP generation, enabling precise control over peptide properties. In this work, we introduce AMP-Diffusion, a latent diffusion model developed to generate functionally potent AMP sequences by applying Gaussian noise over the pre-trained ESM-2-650M embedding space^6,27^. We provide extensive ground-truth experimental validation of AMP-Diffusion-generated peptides by evaluating their structure, mechanism of action, cytotoxicity, and efficacy in animal infection models.

## Results and Discussion

### Computational framework

We developed AMP-Diffusion, a latent space diffusion model designed for antimicrobial peptide (AMP) generation^27^. The model generates new functional AMPs by applying a diffusion process in the latent space of ESM-2 embeddings, progressively adding and then removing noise to generate novel peptide sequences. AMP-Diffusion was trained on a compiled dataset of AMPs from DRAMP 3.0^28^, APD3^29^ and DBAASP^30^ databases (**Dataset S1**).

Our evaluation shows that AMP-Diffusion-generated peptides closely match experimentally validated AMPs in terms of pseudo-perplexity and amino acid residue diversity. These generated sequences also display physicochemical properties comparable to those of naturally occurring AMPs, suggesting their biological plausibility and potential for empirical validation^27^. Model derivation and details can be found in Chen, et al.^27^ For this study, we hypothesized that our pre-trained AMP-Diffusion model would enable the generation of functionally potent AMPs.

From an initial set of 50,000 AMP candidates (**Dataset S2**), we selected 46 peptides for experimental validation based on three criteria: high predicted antimicrobial activity, low similarity to existing AMPs, and broad sequence diversity (**Fig. 1a**, see **Computational Filtering** in the **Methods** section for the peptide selection criteria details). The MIC distribution of our generated sequences was comparable to the training set, indicating the model’s ability to recapitulate the MIC profile of training AMPs (**Fig. 1b** and **S1**). To assess the naturalness of the generated peptides, we employed ProGen2, a protein language model trained on millions of natural protein sequences. Naturalness was quantified using Perplexity scores, where lower values indicate greater naturalness. Our generated sequences exhibited perplexity values (17.90; Filtered 15.83) similar to those of the training set (17.93, two sided Mann-Whitney U-test P-value: 0.0357) (**Fig. 1c**), suggesting that AMP-Diffusion produces peptides with a degree of naturalness comparable to naturally occurring AMPs.

**Figure 1.**
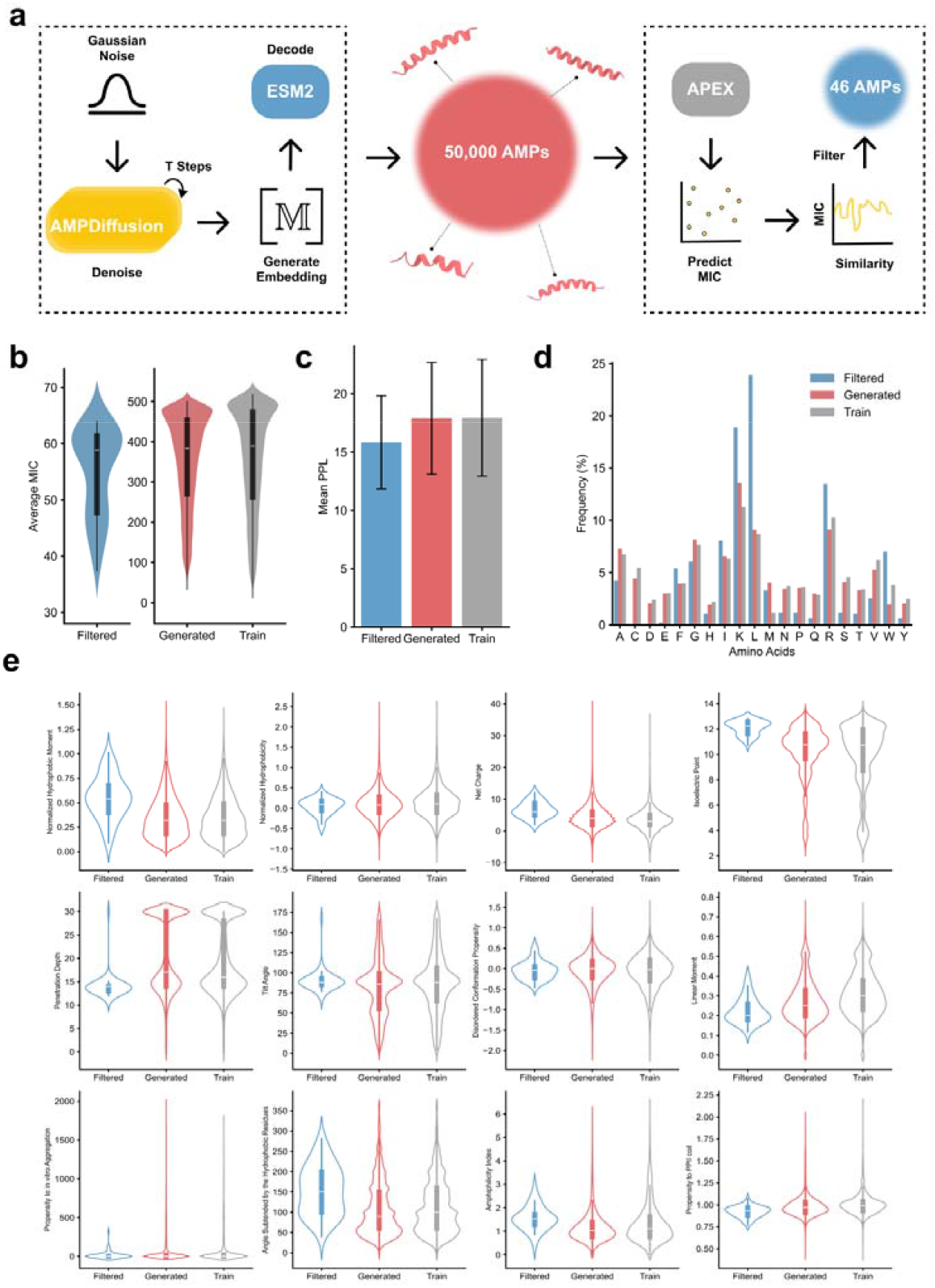
**(a)** Generation and filtration process: 50,000 sequences were generated from random Gaussian noise using the AMP-Diffusion model^27^. Generated Embedding Matrix M is decoded into sequence via ESM2-650M Language Model Head. All sequences underwent MIC prediction for 11 species using the APEX model. A filtered set of 46 peptides was constructed based on average MIC values and sequence identity criteria. **(b)** Distribution of average Minimum Inhibitory Concentration (MIC) values across filtered, generated, and training sets, presented as a bar plot. **(c)** Comparison of perplexity (PPL) scores among the three groups (filtered, generated, and training sets), visualized as a bar plot. **(d)** Amino acid distribution profiles for the filtered, generated, and training sets. **(e)** Distribution of key physicochemical features across the three groups.

### Composition and physicochemical features of the predicted peptides

The generated AMP library closely resembled the training set in amino acid composition, indicating that our model successfully captured the compositional characteristics of known AMPs (**Fig. 1d**). Interestingly, the filtered subset of generated sequences exhibited a distinct pattern, with notably higher frequencies of lysine (K), leucine (L), and arginine (R). These residues are known to be important for initial electrostatic interactions (K and R) with negatively charged bacterial membranes and for amphiphilicity (K, L, and R), which facilitates effective lipid membrane interactions ^11,31,32^. Thus, enrichment in cationic and aliphatic residues directly influences the physicochemical features of the AMP-Diffusion-predicted peptides (**Fig. 1e**), making the filtered set slightly more amphiphilic and highly charged, with enhanced structure tendencies and increased antimicrobial activity.

### *In vitro* antimicrobial activity

To validate our model’s predictive accuracy, we synthesized and experimentally tested 46 peptide sequences. Selection for synthesis was guided by predicted antimicrobial activity and sequence diversity, ensuring broad representation of the generated peptide space (see **APEX Model** and **Computational Filtering** in the **Methods** section).

Thirty five (76%) out of 46 synthesized peptides were active against at least one of the 11 clinically relevant bacterial pathogens tested, including members of the ESKAPEE pathogen list^1^ (**Fig. 2a**). Most peptides exhibited broad-spectrum activity, with *A. baumannii* ATCC 19606 and vancomycin-resistant *E. faecium* ATCC 700221 emerging as the most susceptible strains.

**Figure 2.**
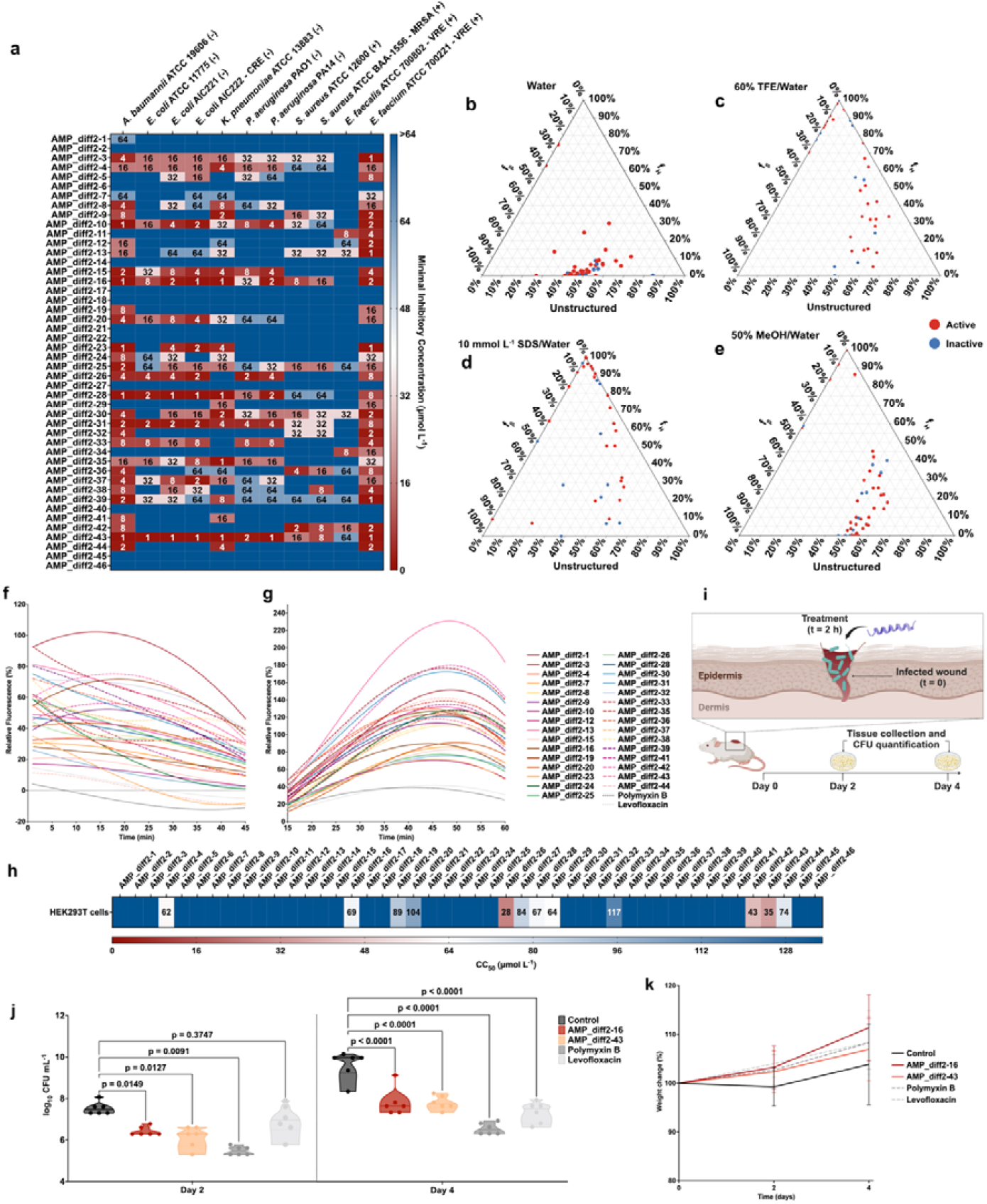
Antimicrobial activity, secondary structure, mechanism of action, cytotoxicity and anti-infective activity in a murine infection model of the peptides generated with AMP-Diffusion. **(a)** Heat map of the antimicrobial activities (μmol L^-1^) of the synthesized peptides against 11 clinically relevant pathogens, including four strains resistant to conventional antibiotics. Briefly, 10^6^ bacterial cells and serially diluted peptides (1-64 μmol L^-1^) were incubated at 37 °C. One day post-treatment, the optical density at 600 nm was measured in a microplate reader to evaluate bacterial growth in the presence of the peptides. MIC values in the heat map are the mode of the three replicates in each condition. **(b-e)** Ternary plots showing the percentage of secondary structure for each peptide (at 50 μmol L^-1^) in four different solvents: **(b)** water, 60% **(c)** trifluoroethanol (TFE) in water, **(d)** Sodium dodecyl sulfate (SDS, 10 mmol L^-1^) in water, and **(e)** 50% methanol (MeOH) in water. Secondary structure fractions were calculated using the BeStSel server^47^. Red dots indicate active peptides, while blue dots represent inactive peptides (see also **Fig. S2**). **(f-g)**To assess whether the peptides act on bacterial membranes, all active peptides against *A. baumannii* ATCC 19606 were subjected to outer membrane permeabilization and cytoplasmic membrane depolarization assays. The fluorescent probe 1-(N-phenylamino)naphthalene (NPN) was used to assess membrane permeabilization **(f)** induced by the tested peptides. The fluorescent probe 3,3′-dipropylthiadicarbocyanine iodide (DiSC_3_-5) was used to evaluate membrane depolarization **(g)** caused by the peptides. The values displayed represent the relative fluorescence of both probes, with non-linear fitting compared to the baseline of the untreated control (buffer + bacteria + fluorescent dye) and benchmarked against the antibiotics polymyxin B and levofloxacin (see also **Fig. S3**). **(h)** Cytotoxic concentrations leading to 50% cell lysis (CC_50_) were determined by interpolating the dose-response data using a non-linear regression curve. All experiments were performed in three independent replicates. **(i)** Schematic of the skin abscess mouse model used to assess selected peptides (n□=□6) for activity against *A. baumannii* ATCC 19606. **(j)** The peptides AMP-diff2-16 and 43, administered at 10 μmol L^-1^ in a single dose, significantly inhibited bacterial proliferation for up to four days post-treatment compared to the untreated control group. Both peptides reduced the infection to a level comparable with the antibiotic controls polymyxin B and levofloxacin. **(k)** Mouse weight was monitored throughout the duration of the skin abscess model (4 days total) to assess potential toxicity from bacterial load or peptide treatment. Statistical significance in **j** was determined using one-way ANOVA followed by Dunnett’s test; p values are shown in the graph. Figure panel **i** was created with BioRender.com.

### Secondary structure

AMPs often adopt an α-helical conformation upon interaction with bacterial membranes^11^. This structural transition is influenced by amphipathicity, net charge, hydrophobicity, and peptide length. To assess secondary structure under various biochemical environments, we tested each peptide in water (extracellular environment), a helix-inducing medium (trifluoroethanol in water, 3:2, v/v), a β-inducing medium (methanol in water, 1:1, v/v), and a micelle solution (10 mmol L□^1^ SDS) simulating lipid bilayer interactions.

Consistent with their high lysine (K), leucine (L), and arginine (R) content and in line with known AMPs, the peptides underwent a helix-coil transition. They adopted α-helical conformations in helix-inducing medium and lipid bilayer environments, while remaining unstructured in water and methanol/water mixtures (**Fig. 2b-e and Fig. S2**).

### Mechanism of action

The bacterial membrane is a common target for most known AMPs due to their non-specific interactions with the lipid bilayer^11^. The antimicrobial activity of AMPs is influenced by their amino acid composition, distribution, and physicochemical characteristics such as amphiphilicity and hydrophobicity.

To interrogate the bactericidal mechanism, we first tested whether they permeabilized the outer membrane of *A. baumannii* using 1-(N-phenylamino)naphthalene (NPN) assays. NPN fluoresces more intensely in lipidic than in aqueous environments, allowing the assessment of membrane integrity. All tested peptides were at least as active as polymyxin B and levofloxacin (antibiotic controls). The peptides AMP-diff2-1, -13, -32, -33, and -38 showed particularly high levels of outer membrane permeabilization (**Fig. 2f** and **S3a**). These results indicate that the peptides are more effective at permeabilizing bacterial membranes than many previously reported AMPs and encrypted peptides (EPs) derived from human proteins, extinct organisms, or bacterial metagenomes^16,18,26,34^. Outer membrane permeabilization is the most common mechanism of action described for AMPs.

Next, we evaluated whether the peptides depolarized the cytoplasmic membrane of *A. baumannii*. Using the potentiometric fluorophore DiSC_3_-5, which fluoresces upon membrane depolarization, we found that the peptides caused stronger depolarization than polymyxin B and levofloxacin. Their activity was comparable to EPs known for their potent depolarizer properties^18^. AMP-diff2-13, 30, 32, 33, and 42 were the most potent depolarizers (**Fig. 2g** and **S3b**).

### Cytotoxicity assays

All 46 synthesized peptides were tested for cytotoxic activity against human embryonic kidney (HEK293T) cells, a well-characterized line commonly used for toxicity^31,36,38^. Among these, 34 displayed no significant cytotoxicity at the tested concentrations (8-128 μmol L^-1^). Cytotoxicity was detected for 12 peptides, and only 6 showed cytotoxicity at concentrations below 64 μmol L^-1^ (the maximum concentration tested in bacterial inhibition assays). These findings reinforce the overall excellent safety profiles of this novel peptide class. The CC_50_ (peptide dose leading to 50% cytotoxicity) was estimated by non-linear regression (**Fig. 2h**).

### Anti-infective efficacy in an animal model

To assess whether the active peptides had anti-infective efficacy *in vivo*, we used preclinical mouse model of skin abscess infection^40,42,44–46^. Two of the peptides, AMP-diff2-16 and 43, with high activity against *A. baumannii* ATCC 19606 (MIC *in vitro* of 1 μmol L^-1^) were tested at a single dose at 10-times MIC concentration (10 μmol L^-1^) after infection was established.

In this model, mice were infected subcutaneously with *A. baumannii* cells (**Fig. 2i**). A single dose of each peptide was administered directly to the infected area. After two days, both peptides markedly reduced the bacterial load by 1.5 orders of magnitude (Fig. 4b). After four days, bacterial counts had decreased by 2–2.5 orders of magnitude, comparable to reductions achieved by widely used antibiotics polymyxin B and levofloxacin^4,8^, which were used as antimicrobial controls (**Fig. 2j**). No weight changes, skin damage, or other adverse effects were observed in treated mice (**Fig. 2k**).

These substantial *in vivo* results highlight the potential of AMP-Diffusion–generated peptides as effective anti-infective agents, on par with standard-of-care antibiotics under physiologically relevant conditions.

### Limitations of the study

The current version of AMP-Diffusion is based on the original Denoising Diffusion Probabilistic Models (DDPM) framework^1^, which has not been optimized for generation quality and computational efficiency compared to more recent methods like Elucidated Diffusion Models (EDM^33^ and EDM2^35^). In addition, AMP-Diffusion operates as an unconditional diffusion model, lacking conditional guidance mechanisms. Future work may explore training a dedicated classifier for noised sequences to guide generation based on inhibitory activity or, alternatively, using classifier-free guidance with pathogen-specific embeddings to produce peptides tailored for particular bacterial strains. Finally, we believe that novel discrete diffusion architectures, such as Masked Discrete Diffusion (MDLM) and Discrete Denoising Posterior Prediction (DDPP), will be apt architectural candidates for a next-generation AMP-Diffusion model^37,39,41^. Incorporating these innovations could yield a next-generation framework that further refines peptide design for enhanced antimicrobial efficacy and safety.

## Methods

### AMP-Diffusion

AMP-Diffusion is a latent space diffusion model designed for generating antimicrobial peptides (AMPs) following DDPM framework. It leverages the state-of-the-art ESM-2 protein language model to map peptide sequences into a continuous latent space, where Gaussian noise is systematically added during the forward process. The model then trains a denoiser to reconstruct the original latent embeddings from the noised inputs, minimizing the l2 loss between predicted and original latent embeddings. For generation, the reverse process starts with Gaussian noise and iteratively denoises it to produce samples resembling the original data distribution. The denoising architecture employs pre-trained ESM-2-8M attention blocks integrated with positional time embeddings and a multilayer perceptron (MLP) for final processing. For the purposes of this project, we retrained the model on a clean and expanded dataset (size = 19,670 peptides, **Dataset S1**) from DRAMP, DBAASP and APD3 using Exponential Moving Average. All other parameters and methodologies remained consistent with the original model architecture. For a comprehensive understanding of the technical details, it is encouraged to refer to the original research.

### APEX Model

We utilized APEX^18^, a bacterial strain-specific antimicrobial activity predictor for peptide sequences to rank and select the generated peptides (https://gitlab.com/machine-biology-group-public/apex-pathogen). As APEX predicts antimicrobial activities against eleven pathogen strains, we used the mean prediction (minimum inhibitory concentration, MIC) to sort the peptides obtained by AMP-Diffusion. Top-scored peptides that were not filtered out by our diversity and similarity criteria (see **Computational Filtering** section) were selected for synthesis and validation.

### Computational Filtering

To ensure that the selected sequences for validation were (a) showing high antimicrobial activities, (b) sufficiently distinct from known AMPs and (c) covered a diverse sequence space, we applied three filtering criteria to the generated peptide sequences: (1) the generated peptides must have mean MIC ≤ 64 μmol L^-1^. (2) similarity to known AMPs, where we calculated sequence similarity for each generated peptide using local sequence alignment^18^ against a curated set of AMPs from DRAMP 3.0^28^, DBAASP^30^, and APD3^29^, along with peptides from our inventory. Peptides with a sequence similarity > 0.60 to any known AMP were considered too similar and were excluded from further analysis. (3) diversity of remaining peptides, in which, for the remaining peptides, we performed pairwise sequence similarity comparisons. When two peptides had a sequence similarity > 0.40, we retained only the one with the lower predicted mean MIC, as estimated by APEX^18^.

### ProGen2 and Perplexity

ProGen2 is a suite of protein language models developed for protein sequence generation and understanding. The model is based on the autoregressive language model. ProGen2 comes in various sizes, ranging from 151 million to 6.4 billion parameters, allowing for different computational requirements and applications. The models use a standard transformer decoder with left-to-right causal masking and incorporate rotary positional encodings. ProGen2 was trained on a mixture of protein sequences from diverse sources including UniRef90, UniProtKB, and BFD30. This diverse training data allows ProGen2 to learn patterns and relationships across a wide range of protein families and structures.

Perplexity measures how well a probabilistic model predicts a sample and is commonly used in language models, including protein language models, to evaluate sequence quality. The perplexity is defined as the exponential of the average negative log-likelihood per token.

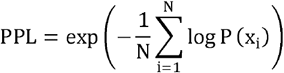

where N is the number of tokens, and P(x_i_) is the predicted probability of the i-th token. Lower perplexity indicates better model confidence. For protein language models like ProGen2, perplexity is associated with the model’s ability to capture the distribution of observed evolutionary sequences. In our study, we utilized the ProGen2-medium for scoring. Lower perplexity suggests a higher likelihood of the model producing valid and biologically relevant protein structures, reflecting its capability to generate novel, viable protein sequences and predict protein fitness without additional fine-tuning.

### Physicochemical Features

Generated peptides from AMP-Diffusion were compared with natural AMPs (the training data). The physicochemical properties, including Normalized Hydrophobic Moment, Normalized Hydrophobicity, Net Charge, Isoelectric Point, Penetration Depth, Tilt Angle, Disordered Conformation Propensity, Linear Moment, Propensity to in vitro Aggregation, Angle Subtended by the Hydrophobic Residues, Amphiphilicity Index, and Propensity to PPII coil, were calculated using the DBAASP server^30^.

### Peptide Synthesis

All peptides for the experiments were obtained from AAPPTec and synthesized using solid-phase peptide synthesis with the Fmoc strategy.

### Minimal inhibitory concentration assays

Broth microdilution assays^43^ were performed to determine the minimum inhibitory concentration (MIC) of each peptide. Peptides were first added to untreated polystyrene 96-well microtiter plates and then serially diluted two-fold in sterile water, with final concentrations ranging from 1 to 64 μmol L^-1^. A bacterial inoculum, prepared at a density of 10^6^ CFU mL^-1^ in LB medium, was mixed with the peptide solutions at a 1:1 ratio. Following a 24-hour incubation at 37 °C, the MIC was defined as the lowest peptide concentration that completely inhibited bacterial growth. Each assay was conducted in triplicate across three independent experiments.

### Circular dichroism experiments

Circular dichroism (CD) experiments were carried out using a J1500 circular dichroism spectropolarimeter (Jasco) at the Biological Chemistry Resource Center (BCRC) of the University of Pennsylvania. Measurements were conducted at 25 °C, with spectra representing the average of three accumulations. A quartz cuvette with a 1.0 mm optical path length was used, and data were collected from 260 to 190 nm at a scanning rate of 50 nm min^-1^ with a bandwidth of 0.5 nm. Peptides were tested at a concentration of 50 μmol L^-1^ in four different solvent conditions: water, a 3:2 trifluoroethanol (TFE)/water mixture, 10 mmol L^-1^ sodium dodecyl sulfate (SDS) in water, and a 1:1 methanol (MeOH)/water mixture, with corresponding baselines recorded prior to measurement. A Fourier transform filter was applied to reduce background noise. Secondary structure fractions were determined using the BeStSel^47^ single spectra analysis tool. Ternary plots were generated using https://www.ternaryplot.com/ and subsequently modified for visualization.

### Outer membrane permeabilization assays

The N-phenyl-1-naphthylamine (NPN) uptake assay was employed to assess the ability of peptides to permeabilize the bacterial outer membrane. Cultures of *A. baumannii* ATCC 19606 were grown to an optical density at 600 nm (OD600) of 0.4, centrifuged at 10,000 rpm for 10 minutes at 4 °C, washed, and resuspended in 5 mmol L^-1^ HEPES buffer (pH 7.4) supplemented with 5 mmol L^-1^ glucose. The bacterial suspension (100 μL) was added to a white 96-well plate, followed by the addition of 4 μL of NPN at a final concentration of 0.5 mmol L□^1^. Peptides diluted in water were subsequently added to each well, and fluorescence measurements were recorded over 45 minutes at an excitation wavelength (λ_ex_) of 350 nm and an emission wavelength (λ_em_) of 420 nm. Relative fluorescence was calculated using an untreated control (buffer + bacteria + fluorescent dye) as the baseline, with the percentage difference between the baseline and sample values determined using the following equation:

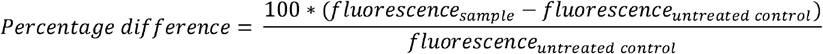

### Cytoplasmic membrane depolarization assays

The cytoplasmic membrane depolarization assay utilized the membrane potential-sensitive dye 3,3 ′ -dipropylthiadicarbocyanine iodide (DiSC_3_-5) to evaluate peptide activity. Mid-logarithmic phase cultures of *A. baumannii* ATCC 19606 were washed and resuspended in HEPES buffer (pH 7.2) containing 20 mmol L^-1^ glucose and 0.1 mol L^-1^ KCl at a concentration of 0.05 OD_600_ mL^-1^. DiSC_3_-5 was added to the bacterial suspension at a final concentration of 20 μmol L^-1^ and incubated for 15 minutes to allow stabilization of fluorescence, indicating dye incorporation into the bacterial membrane. Peptides were then mixed with the bacterial suspension at a 1:1 ratio to achieve final concentrations equivalent to their MIC values. Membrane depolarization was monitored by measuring fluorescence changes (λ_ex_ = 622 nm, λ_em_ = 670 nm) over 60 minutes. Relative fluorescence was calculated using an untreated control (buffer + bacteria + dye) as the baseline, and the percentage difference between the baseline and the sample was determined using the following equation:

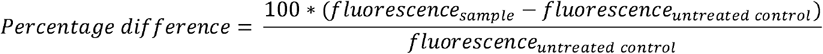

### Eukaryotic cell culture conditions and cytotoxicity assays

HEK293T cells were obtained from the American Type Culture Collection (CRL-3216). The cells were cultured in high-glucose Dulbecco’s modified Eagle’s medium supplemented with 1% penicillin and streptomycin (antibiotics) and 10% fetal bovine serum and grown at 37□°C in a humidified atmosphere containing 5% CO_2_.

One day before the experiment, an aliquot of 100□μL of the cells at 50,000□cells per mL was seeded into each well of the cell-treated 96-well plates used in the experiment (that is, 5,000 cells per well). The attached HEK293T cells were then exposed to increasing concentrations of the peptides (8–128□μmol□L^-1^) for 24□h. After the incubation period, we performed the 3-(4,5-dimethylthiazol-2-yl)-2,5-diphenyltetrazolium bromide tetrazolium reduction assay (MTT assay). The MTT reagent was dissolved at 0.5□mg□mL^-1^ in medium without phenol red and was used to replace cell culture supernatants containing the peptides (100□μL per well), and the samples were incubated for 4□h at 37□°C in a humidified atmosphere containing 5% CO_2_ yielding the insoluble formazan salt. The resulting salts were then resuspended in hydrochloric acid (0.04□mol□L^-1^) in anhydrous isopropanol and quantified by spectrophotometric measurements of absorbance at 570□nm. All assays were done as three biological replicates.

### Skin abscess infection mouse model

Six-week-old female CD-1 mice were anesthetized, and their backs were shaved. The area was then wiped with ethanol to eliminate any endogenous bacteria prior to creating a surface abrasion using a scalpel. An aliquot of *A. baumannii* ATCC 19606 (10^6^ CFU mL^-1^; 20 μL), grown to an optical density of 0.5 at 600 nm in LB medium and washed twice with sterile PBS (pH 7.4) by centrifugation (5,000×g for 3 minutes), was applied to the scratched region. Peptides, polymyxin B or levofloxacin, each diluted to the 10×MIC, was administered to the wound site 2 hours post-infection. Two and four days after the infection, the animals were euthanized, and the infected skin was excised. The tissues were homogenized using a bead beater at 25 Hz for 20 minutes. The samples were then serially diluted 10-fold and plated on MacConkey agar plates for colony-forming unit (CFU) quantification. Six mice per group were included in the study. All procedures were approved by the University Laboratory Animal Resources (ULAR) at the University of Pennsylvania (Protocol 806763).

## Acknowledgments

Cesar de la Fuente-Nunez holds a Presidential Professorship at the University of Pennsylvania and acknowledges funding from the Procter & Gamble Company, United Therapeutics, a BBRF Young Investigator Grant, the Nemirovsky Prize, Penn Health-Tech Accelerator Award, Defense Threat Reduction Agency grants HDTRA11810041 and HDTRA1-23-1-0001, and the Dean’s Innovation Fund from the Perelman School of Medicine at the University of Pennsylvania. Research reported in this publication was supported by the Langer Prize (AIChE Foundation), the NIH R35GM138201, and DTRA HDTRA1-21-1-0014. We thank Dr. Mark Goulian for kindly donating the following strains: *Escherichia coli* AIC221 [*Escherichia coli* MG1655 phnE_2::FRT (control strain for AIC 222)] and *Escherichia coli* AIC222 [*Escherichia coli* MG1655 pmrA53 phnE_2::FRT (polymyxin resistant)]. We thank de la Fuente Lab members for insightful discussions. We also thank Pranay Vure and Rishab Pulugurta for helping model evaluation in previous development round. Figures created with BioRender.com are attributed as such. Molecules were rendered using the PyMOL Molecular Graphics System, Version 3.0 Schrödinger, LLC.

## Author contributions

TC and PC conceived, designed, and developed AMP-Diffusion architecture, training, and analysis. PC directed and supervised the development of AMP-Diffusion. MDTT and CFN conceived and designed the experimental validation of peptides generated with AMP-Diffusion. MDTT performed all experiments. FW performed the computational filtering, physicochemical feature calculation, and peptide visualization. CFN supervised the experiments. MDTT, TC, and FW performed data analysis. CFN and PC funded the project. All authors wrote the original draft and revised the final version.

## Data and materials availability

All training data, testing data, and code used to develop the machine learning model are freely available on GitLab (https://github.com/programmablebio/amp-diffusion). All data pertaining to the experimental validation of generated peptides are available in the Supplementary Data.

## Supplementary Materials

Figs. S1 to S3

## References

1. Ho, J., Jain, A., and Abbeel, P. (2020). Denoising Diffusion Probabilistic Models. NeurIPSProceedings 33, 1–12.

2. Watson, J.L., Juergens, D., Bennett, N.R., Trippe, B.L., Yim, J., Eisenach, H.E., Ahern, W., Borst, A.J., Ragotte, R.J., Milles, L.F., et al. (2023). De novo design of protein structure and function with RFdiffusion. Nature 620, 1089–1100. 10.1038/s41586-023-06415-8.

3. Corso, G., Hannes, S., Jing, B., Barzilay, R., and Jaakkola, T. (2023). DiffDock: Diffusion Steps, Twists, and Turns for Molecular Docking. ArXiv.

4. Peng, Z., Han, C., Wang, X., Li, D., and Yuan, F. (2023). Generative Diffusion Models for Antibody Design, Docking, and Optimization. bioRxiv, 2023.09.25.559190. 10.1101/2023.09.25.559190.

5. Lin, Z., Akin, H., Rao, R., Hie, B., Zhu, Z., Lu, W., Smetanin, N., Verkuil, R., Kabeli, O., Shmueli, Y., et al. (2023). Evolutionary-scale prediction of atomic-level protein structure with a language model. Science (1979) 379, 1123–1130. 10.1126/science.ade2574.

6. Elnaggar, A., Heinzinger, M., Dallago, C., Rehawi, G., Wang, Y., Jones, L., Gibbs, T., Feher, T., Angerer, C., Steinegger, M., et al. (2022). ProtTrans: Toward Understanding the Language of Life Through Self-Supervised Learning. IEEE Trans Pattern Anal Mach Intell 44, 7112–7127. 10.1109/TPAMI.2021.3095381.

7. Ferruz, N., Schmidt, S., and Höcker, B. (2022). ProtGPT2 is a deep unsupervised language model for protein design. Nat Commun 13, 4348. 10.1038/s41467-022-32007-7.

8. Madani, A., McCann, B., Naik, N., Keskar, N.S., Anand, N., Eguchi, R.R., Huang, P.-S., and Socher, R. (2020). ProGen: Language Modeling for Protein Generation. ArXiv.

9. Alamdari, S., Thakkar, N., van den Berg, R., Tenenholtz, N., Strome, R., Moses, A.M., Lu, A.X., Fusi, N., Amini, A.P., and Yang, K.K. (2023). Protein generation with evolutionary diffusion: sequence is all you need. Preprint, 10.1101/2023.09.11.556673 https://doi.org/10.1101/2023.09.11.556673.

10. Torres, M.D.T., and de la Fuente-Nunez, C. (2019). Reprogramming biological peptides to combat infectious diseases. Chemical Communications 55, 15020–15032. 10.1039/C9CC07898C.

11. Torres, M.D.T., Sothiselvam, S., Lu, T.K., and de la Fuente-Nunez, C. (2019). Peptide Design Principles for Antimicrobial Applications. J Mol Biol 431, 3547–3567. 10.1016/j.jmb.2018.12.015.

12. Fjell, C.D., Hiss, J.A., Hancock, R.E.W., and Schneider, G. (2011). Designing antimicrobial peptides: form follows function. Nat Rev Drug Discov 11. 10.1038/nrd3591.

13. de la Fuente-Nunez, C., Torres, M.D., Mojica, F.J., and Lu, T.K. (2017). Next-generation precision antimicrobials: towards personalized treatment of infectious diseases. Curr Opin Microbiol 37, 95–102. 10.1016/j.mib.2017.05.014.

14. Torres, M.D.T., Cao, J., Franco, O.L., Lu, T.K., and de la Fuente-Nunez, C. (2021). Synthetic Biology and Computer-Based Frameworks for Antimicrobial Peptide Discovery. ACS Nano 15, 2143–2164. 10.1021/acsnano.0c09509.

15. Magana, M., Pushpanathan, M., Santos, A.L., Leanse, L., Fernandez, M., Ioannidis, A., Giulianotti, M.A., Apidianakis, Y., Bradfute, S., Ferguson, A.L., et al. (2020). The value of antimicrobial peptides in the age of resistance. Lancet Infect Dis. 10.1016/S1473-3099(20)30327-3.

16. Maasch, J.R.M.A., Torres, M.D.T., Melo, M.C.R., and de la Fuente-Nunez, C. (2023). Molecular de-extinction of ancient antimicrobial peptides enabled by machine learning. Cell Host Microbe 31, 1260–1274. 10.1016/j.chom.2023.07.001.

17. Torres, M.D.T., and de la Fuente-Nunez, C. (2019). Toward computer-made artificial antibiotics. Curr Opin Microbiol 51, 30–38. 10.1016/j.mib.2019.03.004.

18. Wan, F., Torres, M.D.T., Peng, J., and de la Fuente-Nunez, C. (2024). Deep-learning-enabled antibiotic discovery through molecular de-extinction. Nat Biomed Eng 8, 854–871. 10.1038/s41551-024-01201-x.

19. Cesaro, A., Bagheri, M., Torres, M., Wan, F., and de la Fuente-Nunez, C. (2023). Deep learning tools to accelerate antibiotic discovery. Expert Opin Drug Discov 18, 1245–1257. 10.1080/17460441.2023.2250721.

20. Wong, F., de la Fuente-Nunez, C., and Collins, J.J. (2023). Leveraging artificial intelligence in the fight against infectious diseases. Science (1979) 381, 164–170. 10.1126/science.adh1114.

21. Wan, F., Wong, F., Collins, J.J., and de la Fuente-Nunez, C. (2024). Machine learning for antimicrobial peptide identification and design. Nature Reviews Bioengineering. 10.1038/s44222-024-00152-x.

22. Liu, G., Catacutan, D.B., Rathod, K., Swanson, K., Jin, W., Mohammed, J.C., Chiappino-Pepe, A., Syed, S.A., Fragis, M., Rachwalski, K., et al. (2023). Deep learning-guided discovery of an antibiotic targeting Acinetobacter baumannii. Nat Chem Biol 19, 1342–1350. 10.1038/s41589-023-01349-8.

23. Stokes, J.M., Yang, K., Swanson, K., Jin, W., Cubillos-Ruiz, A., Donghia, N.M., MacNair, C.R., French, S., Carfrae, L.A., Bloom-Ackermann, Z., et al. (2020). A Deep Learning Approach to Antibiotic Discovery. Cell 180, 688–702.e13. 10.1016/j.cell.2020.01.021.

24. Ma, Y., Guo, Z., Xia, B., Zhang, Y., Liu, X., Yu, Y., Tang, N., Tong, X., Wang, M., Ye, X., et al. (2022). Identification of antimicrobial peptides from the human gut microbiome using deep learning. Nat Biotechnol 40, 921–931. 10.1038/s41587-022-01226-0.

25. Li, C., Sutherland, D., Hammond, S.A., Yang, C., Taho, F., Bergman, L., Houston, S., Warren, R.L., Wong, T., Hoang, L.M.N., et al. (2022). AMPlify: attentive deep learning model for discovery of novel antimicrobial peptides effective against WHO priority pathogens. BMC Genomics 23, 77. 10.1186/s12864-022-08310-4.

26. Santos-Júnior, C.D., Torres, M.D.T., Duan, Y., Rodríguez del Río, Á., Schmidt, T.S.B., Chong, H., Fullam, A., Kuhn, M., Zhu, C., Houseman, A., et al. (2024). Discovery of antimicrobial peptides in the global microbiome with machine learning. Cell 187, 3761–3778. 10.1016/j.cell.2024.05.013.

27. Chen, T., Vure, P., Pulugurta, R., and Chatterjee, P. (2024). AMP-Diffusion: Integrating Latent Diffusion with Protein Language Models for Antimicrobial Peptide Generation. bioRxiv, 2024.03.03.583201. 10.1101/2024.03.03.583201.

28. Shi, G., Kang, X., Dong, F., Liu, Y., Zhu, N., Hu, Y., Xu, H., Lao, X., and Zheng, H. (2022). DRAMP 3.0: an enhanced comprehensive data repository of antimicrobial peptides. Nucleic Acids Res 50, D488–D496. 10.1093/nar/gkab651.

29. Wang, G., Li, X., and Wang, Z. (2016). APD3: the antimicrobial peptide database as a tool for research and education. Nucleic Acids Res 44, D1087–D1093.

30. Pirtskhalava, M., Amstrong, A.A., Grigolava, M., Chubinidze, M., Alimbarashvili, E., Vishnepolsky, B., Gabrielian, A., Rosenthal, A., Hurt, D.E., and Tartakovsky, M. (2021). DBAASP v3: database of antimicrobial/cytotoxic activity and structure of peptides as a resource for development of new therapeutics. Nucleic Acids Res 49, D288–D297. 10.1093/nar/gkaa991.

31. Torres, M.D.T., Pedron, C.N., Higashikuni, Y., Kramer, R.M., Cardoso, M.H., Oshiro, K.G.N., Franco, O.L., Silva Junior, P.I., Silva, F.D., Oliveira Junior, V.X., et al. (2018). Structure-function-guided exploration of the antimicrobial peptide polybia-CP identifies activity determinants and generates synthetic therapeutic candidates. Commun Biol 1, 221. 10.1038/s42003-018-0224-2.

32. Tossi, A., Sandri, L., and Giangaspero, A. (2000). Amphipathic, alpha-helical antimicrobial peptides. Biopolymers 55, 4–30. 10.1002/1097-0282(2000)55:1<4::AID-BIP30>3.0.CO;2-M.

33. Karras, T., Aittala, M., and Laine, S. (2022). Elucidating the Design Space of Diffusion-Based Generative Models. ArXiv.

34. Torres, M.D.T., Brooks, E.F., Cesaro, A., Sberro, H., Gill, M.O., Nicolaou, C., Bhatt, A.S., and de la Fuente-Nunez, C. (2024). Mining human microbiomes reveals an untapped source of peptide antibiotics. Cell 187, 5453–5467. 10.1016/j.cell.2024.07.027.

35. Karras, T., Aittala, M., Lehtinen, J., Hellsten, J., Aila, T., and Laine, S. (2023). Analyzing and Improving the Training Dynamics of Diffusion Models. ArXiv.

36. Nim, S., O’Hara, D.M., Corbi-Verge, C., Perez-Riba, A., Fujisawa, K., Kapadia, M., Chau, H., Albanese, F., Pawar, G., De Snoo, M.L., et al. (2023). Disrupting the α-synuclein-ESCRT interaction with a peptide inhibitor mitigates neurodegeneration in preclinical models of Parkinson’s disease. Nat Commun 14, p150. 10.1038/s41467-023-37464-2.

37. Sahoo, S.S., Arriola, M., Schiff, Y., Gokaslan, A., Marroquin, E., Chiu, J.T., Rush, A., and Kuleshov, V. (2024). Simple and Effective Masked Diffusion Language Models. ArXiv.

38. Silva, O.N., Torres, M.D.T., Cao, J., Alves, E.S.F., Rodrigues, L. V., Resende, J.M., Lião, L.M., Porto, W.F., Fensterseifer, I.C.M., Lu, T.K., et al. (2021). Repurposing a peptide toxin from wasp venom into antiinfectives with dual antimicrobial and immunomodulatory properties. Proceedings of the National Academy of Sciences 118, e2025351118. 10.1073/pnas.2025351118.

39. Goel, S., Thoutam, V., Marroquin, E.M., Gokaslan, A., Firouzbakht, A., Vincoff, S., Kuleshov, V., Kratochvil, H.T., and Chatterjee, P. (2024). MeMDLM: De Novo Membrane Protein Design with Masked Discrete Diffusion Protein Language Models.

40. Silveira, G.G.O.S., Torres, M.D.T., Ribeiro, C.F.A., Meneguetti, B.T., Carvalho, C.M.E., de la Fuente-Nunez, C., Franco, O.L., and Cardoso, M.H. (2021). Antibiofilm Peptides: Relevant Preclinical Animal Infection Models and Translational Potential. ACS Pharmacol Transl Sci 4, 55–73. 10.1021/acsptsci.0c00191.

41. Rector-Brooks, J., Hasan, M., Peng, Z., Quinn, Z., Liu, C., Mittal, S., Dziri, N., Bronstein, M., Bengio, Y., Chatterjee, P., et al. (2024). Steering Masked Discrete Diffusion Models via Discrete Denoising Posterior Prediction.

42. Arqué, X., Torres, M.D.T., Patiño, T., Boaro, A., Sánchez, S., and de la Fuente-Nunez, C. (2022). Autonomous Treatment of Bacterial Infections in Vivo Using Antimicrobial Micro- and Nanomotors. ACS Nano 16, 7547–7558. 10.1021/acsnano.1c11013.

43. Wiegand, I., Hilpert, K., and Hancock, R.E.W. (2008). Agar and broth dilution methods to determine the minimal inhibitory concentration (MIC) of antimicrobial substances. Nat Protoc 3, 163–175. 10.1038/nprot.2007.521.

44. Torres, M.D.T., Voskian, S., Brown, P., Liu, A., Lu, T.K., Hatton, T.A.A., and de la Fuente-Nunez, C. (2021). Coatable and Resistance-Proof Ionic Liquid for Pathogen Eradication. ACS Nano 15, 966–978. 10.1021/acsnano.0c07642.

45. Cesaro, A., Torres, M.D.T., Gaglione, R., Dell’Olmo, E., Di Girolamo, R., Bosso, A., Pizzo, E., Haagsman, H.P., Veldhuizen, E.J.A., de la Fuente-Nunez, C., et al. (2022). Synthetic Antibiotic Derived from Sequences Encrypted in a Protein from Human Plasma. ACS Nano 16, 1880–1895. 10.1021/acsnano.1c04496.

46. Torres, M.D.T., Cesaro, A., and de la Fuente-Nunez, C. (2024). Peptides from non-immune proteins target infections through antimicrobial and immunomodulatory properties. Trends Biotechnol. 10.1016/j.tibtech.2024.09.008.

47. Micsonai, A., Moussong, É., Wien, F., Boros, E., Vadászi, H., Murvai, N., Lee, Y.-H., Molnár, T., Réfrégiers, M., Goto, Y., et al. (2022). BeStSel: webserver for secondary structure and fold prediction for protein CD spectroscopy. Nucleic Acids Res 50, W90–W98. 10.1093/nar/gkac345.

