## Supplementary material for "Generative latent diffusion language modeling yields anti-infective synthetic peptides": SI

Supplementary Materials

Figs. S1 to S3

**Supplementary Figures:**

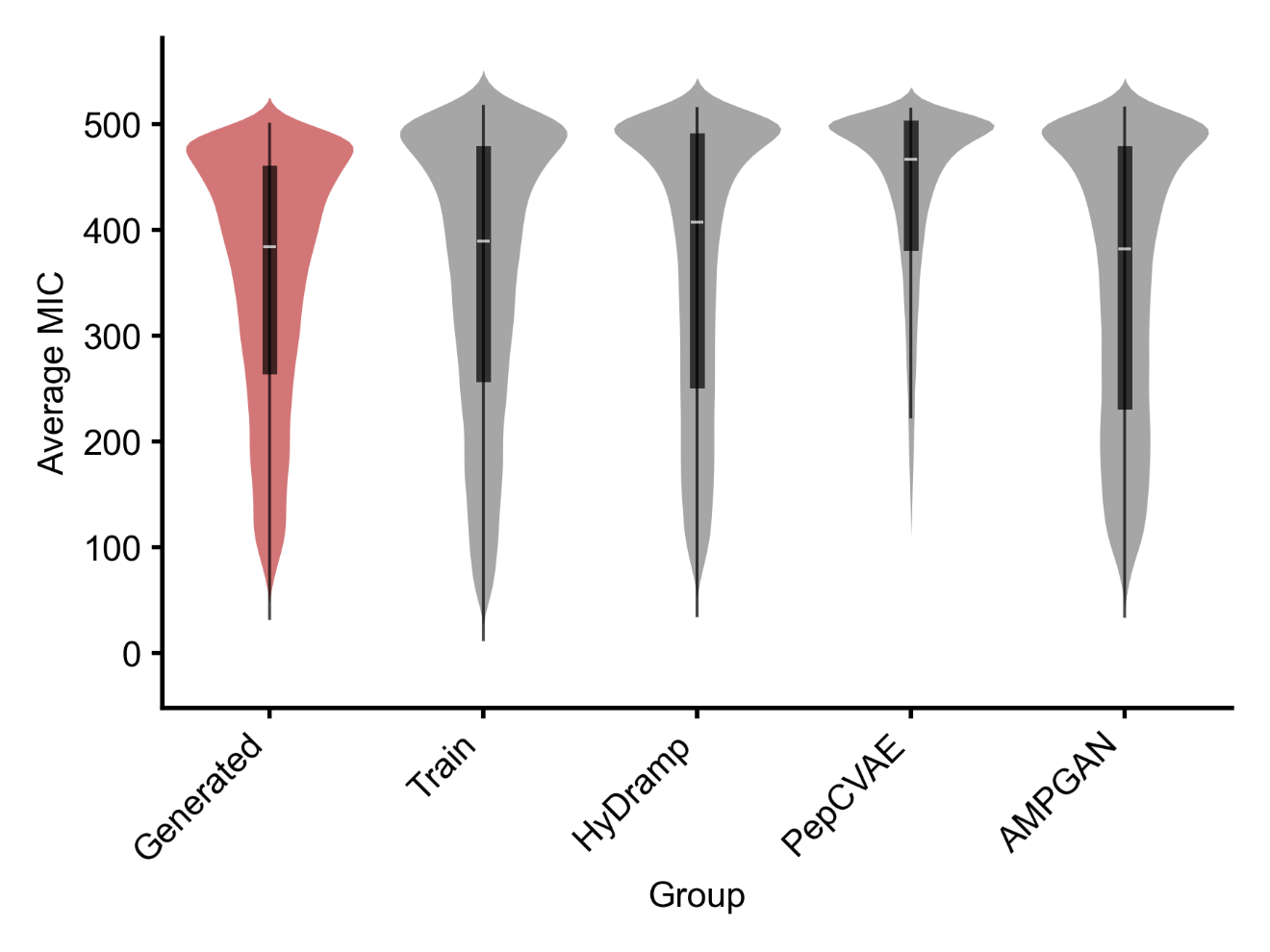

**Figure S1. Comparison of Average Minimum Inhibitory Concentrations Across Different Models.** The analysis compared 50,000 antimicrobial peptides generated by the proposed model with an equal number of peptides generated by HyDramp, PepCVAE, and AMPGAN. The APEX algorithm predicted Minimum Inhibitory Concentration (MIC) values across 11 bacterial species for all generated peptides. (a) The graph displays average predicted MIC values for peptides from the proposed model, training dataset, and comparative models.

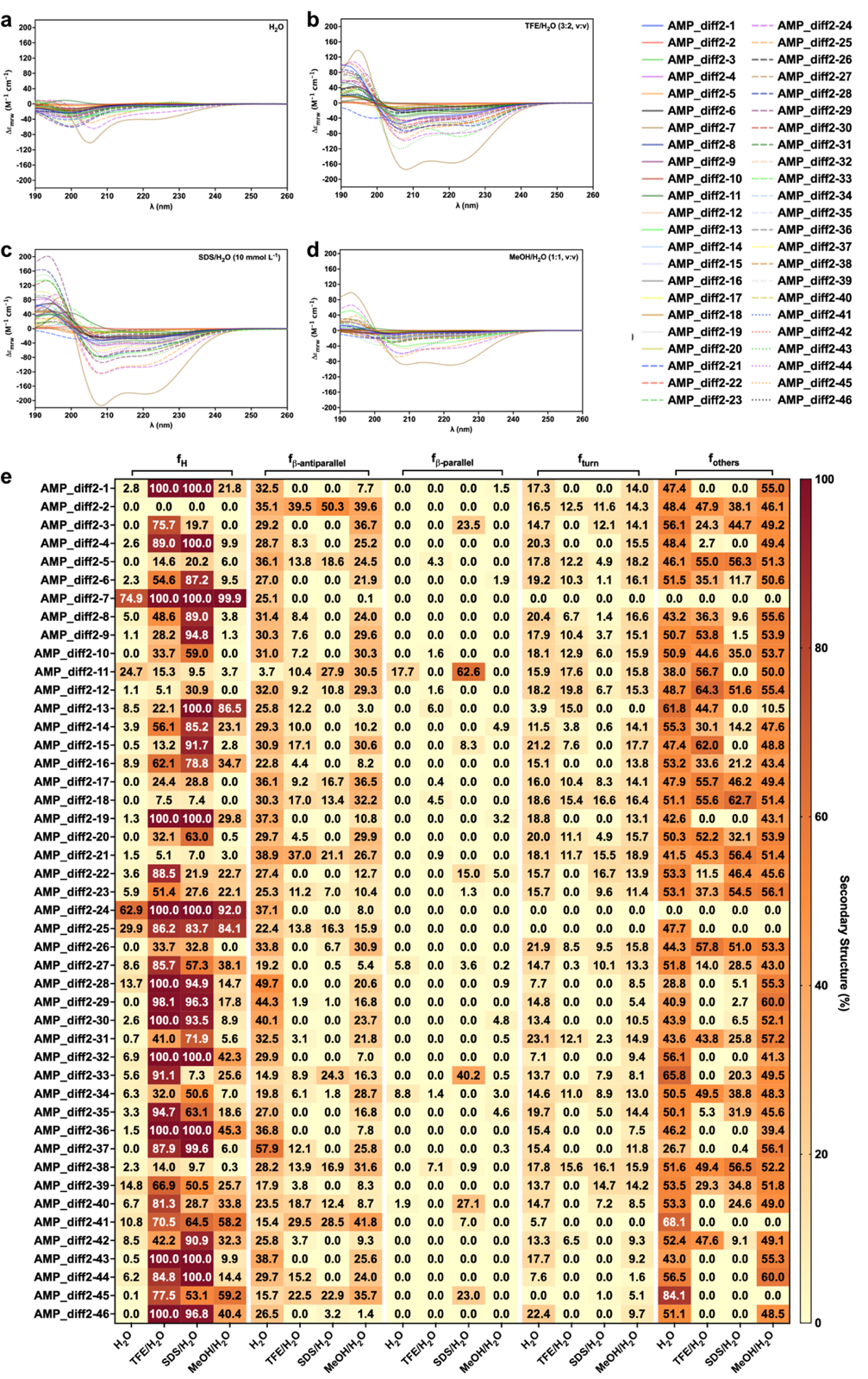

**Figure S2. Circular dichroism spectra of the peptides.** Circular dichroism experiments were conducted with peptides using a J-1500 Jasco circular dichroism spectrophotometer. The spectra were recorded in three different media: **(a)** water, **(b)** 60% trifluoroethanol in water, **(c)** sodium dodecyl sulfate (SDS) in water (10 mmol L^-1^), and (d) 50% methanol in water after three accumulations at 25 ^o^C, using a 1mm path length quartz cell, between 260 and 190 nm at 50 nm min^-1^, with a bandwidth of 0.5 nm. The concentration of all peptides tested was 50 μmol L^-1^. **(d)** Heatmap with the percentage of secondary structure found for each peptide in three different solvents: water, 60% trifluoroethanol in water, Sodium dodecyl sulfate (10 mmol L^-1^) in water, and 50% methanol in water. Secondary structure fraction was calculated using the BeStSel server^15^.

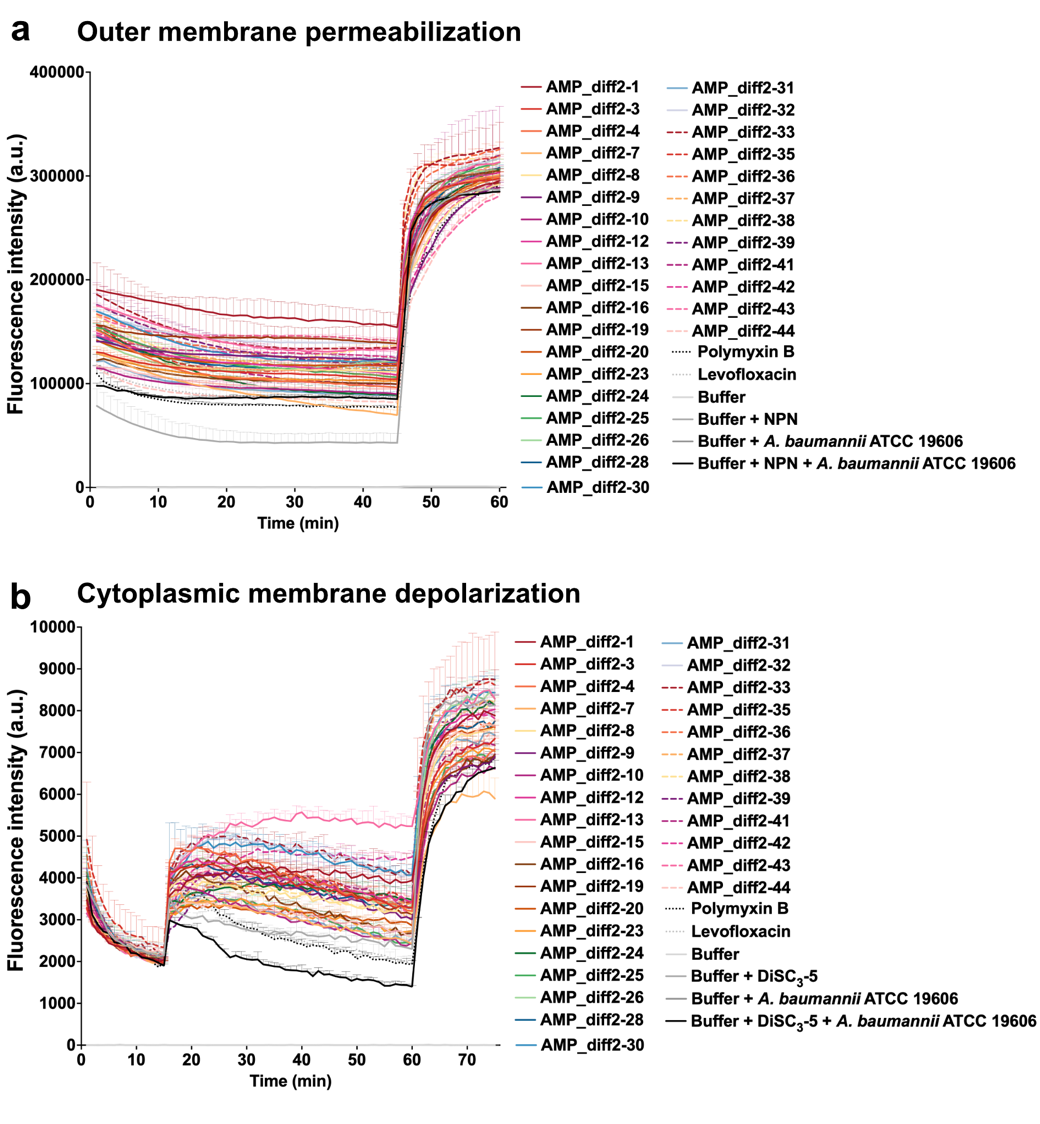

**Figure S3.** **Outer membrane permeabilization and cytoplasmic membrane depolarization of *A. baumannii* ATCC 19606 induced by the peptides. (a)** Outer membrane permeabilization was assessed using the probe 1-(N-phenylamino)naphthalene (NPN), showing the permeabilization effects of the peptides active against *A. baumannii* ATCC 19606. **(b)** Membrane depolarization assays were performed using the hydrophobic probe 3,3′-dipropylthiadicarbocyanine iodide [DiSC_3_-(5)] on all peptides active against *A. baumannii* ATCC 19606. Polymyxin B and levofloxacin served as positive controls, while buffer, buffer with the probe, and buffer with both probe and bacteria were used as baseline controls for fluorescence. The panels display the raw fluorescence intensity data obtained from the experiments. Error bars are the standard deviation obtained from the three replicates.
